# The impact of mutation L138F/L210F on the Orai channel: a molecular dynamics simulation study

**DOI:** 10.1101/2021.06.23.449681

**Authors:** Xiaoqian Zhang, Hua Yu, Xiangdong Liu, Chen Song

## Abstract

The calcium release-activated calcium (CRAC) channel, composed of the Orai channel and the STIM protein, plays a crucial role in maintaining the Ca^2+^ concentration in cells. Previous studies showed that the L138F mutation in the human Orai1 creates a constitutively open channel independent of STIM, causing severe myopathy, but how the L138F mutation activates Orai1 is still unclear. Here, based on the crystal structure of *Drosophila melanogaster* Orai (dOrai), molecular dynamics simulations for the wild-type (WT) and the L210F (corresponding to L138F in the human Orai1) mutant were conducted to investigate their structural and dynamical properties. The results showed that the L210F dOrai mutant tends to have a more hydrated hydrophobic region (V174 to F171), as well as more dilated basic region (K163 to R155) and selectivity filter (E178). Sodium ions were located deeper in the mutant than in the WT. Further analysis revealed two local but essential conformational changes that may be the key to the activation. A rotation of F210, a previously undescribed feature, was found to result in the opening of the K163 gate through hydrophobic interactions. At the same time, a counter-clockwise rotation of F171 occurred more frequently in the mutant, resulting in a wider hydrophobic gate with more hydration. Ultimately, the opening of the two gates may facilitate the opening of the Orai channel independent of STIM.

## Introduction

Calcium ions, as an essential second messenger in cells, regulate a wide range of physiological processes. Store-operated calcium entry (SOCE) was identified to explain how depletion of endoplasmic reticulum (ER) Ca^2+^ stores evokes Ca^2+^ influx across the plasma membrane.^1^ Up to now, the relatively well-studied SOCE channel is the “calcium release-activated calcium” (CRAC) channel, which is involved in numerous cell activities such as gene transcription, muscle contraction, secretion, cell proliferation, differentiation and apoptosis etc.^2–5^ Both loss-of-function and gain-of-function mutations of the CRAC channel lead to devastating immunodeficiencies, bleeding disorders and muscle weakness.^6–9^ In recent decades, our understanding of the operational mechanisms of the CRAC channel including the gating mechanism has been greatly advanced, with the discovery of its molecular components, stromal interaction molecule (STIM) and the pore-forming protein Orai.^10, 11^ The STIMs are single-pass ER transmembrane proteins, function as the sensor of the Ca^2+^ concentration inside the ER, bind to and activate Orai channels.^11, 12^ Two mammalian homologs, STIM1 and STIM2, are included in the STIMs family and the former one is more widely studied. Orai, the calcium channel that opens to permit the influx of the calcium ions, locates on the plasma membrane and contains three closely conserved mammalian homologs, Orai1, Orai2 and Orai3.^13, 14^

Orai1 has a high calcium selectivity (> 1000-fold over Na^+^) and low conductivity (< 1 pS).^15, 16^ According to the structure of Orai (Fig. 1A),^17–19^ the transmembrane Orai is composed of six subunits with a central pore formed by six helices denoted as transmembrane one (TM1). TM1 are surrounded by two rings: one is composed of TM2 and TM3, the other is TM4. There is another helix which extends into the cytosol, termed TM4 extension. As TM1 helices are tightly wrapped by TM2 and TM3 helices, they may have limited space to expand to allow the CRAC channel open.^11, 20^ The TM1 helices can be divided into four distinct regions (Fig. 1A): the selectivity filter (SF) - a ring of glutamates (E178), the hydrophobic region (V174, F171, L167), the basic region (K163, K159, R155) and the cytosolic region.^17^ The glutamate-ring (E178) functions as a SF and makes the channel have a high calcium ion selectivity, which is the most significant feature of Orai channels. Mutation of the residue E178 to aspartate disrupts Ca^2+^-selectivity.^22^ The well-packed side chains of V174, F171 and L167 form the inner wall of the hydrophobic region, having extensive hydrophobic interactions with one another, and are strictly conserved among Orai channels.^23, 24^ These hydrophobic residues are located at the center of the protein, which likely form a gate of the pore. The V174A mutation yields an activated channel with altered ion selectivity even if its pore structure shows no obvious changes compared to the wild-type (WT), and a slight difference of the number of water molecules in the hydrophobic region is enough to change the conduction state of the pore,^25^ indicating the significant role of the hydrophobic region in gating. Another important region locates in the lower part of the channel and lines by three basic residues (K163, K159 and R155), creating an unexpected positively charged environment for the pore that conducts cations. Generally, K163 corresponds to the narrowest point of the pore, resulting in large electrostatic repulsion between this positively charged residue and cations passing by. Therefore, K163 is believed to be the other gate of the pore and jointly regulates the channel state together with the hydrophobic gate.^26^

**Figure 1.**
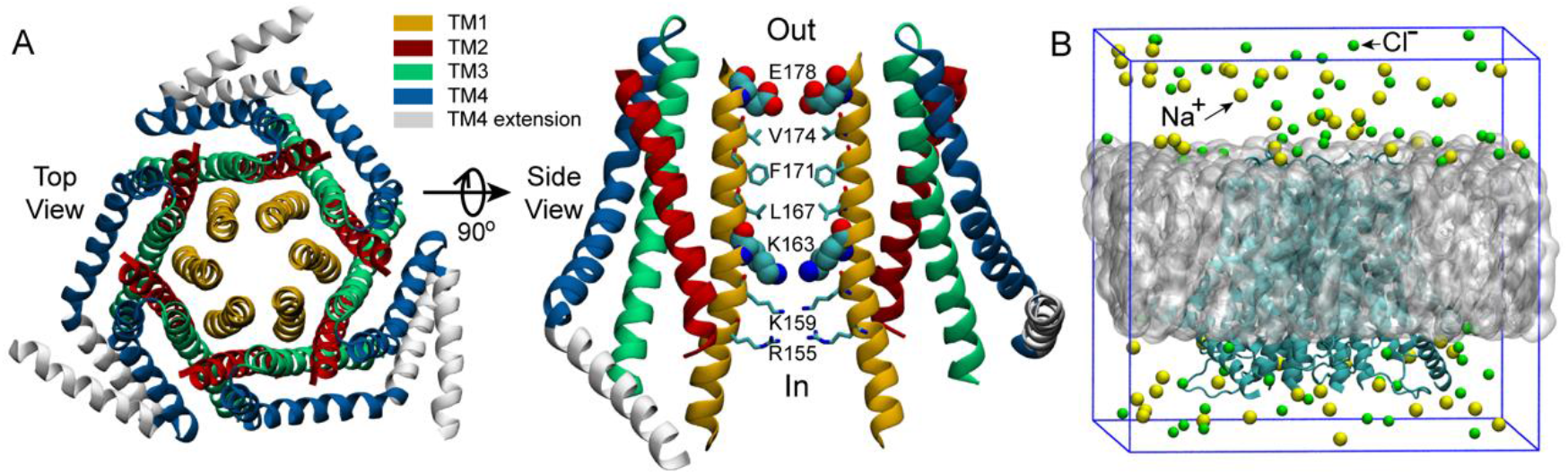
The structure of dOrai and the initial simulation system. (A) The crystal structure of *Drosophila melanogaster* Orai obtained from the Protein Data Bank (PDB ID: 4HKR)^17^. In the right panel, only two oppositing subunits are shown for clarity. The SF of the pore, E178, and the starting residue of the basic region, K163, are shown with the VDW representation, the other pore residues are shown with Licorice representation in VMD (visual molecular dynamics)^21^. (B) The initial simulation system. The grey sphere represents the POPC bilayer. Water molecules are not shown here but included in the simulations.

Previous experiments have shown that the L138F mutation in human Orai1 yields a constitutively permeant channel that allows ion conduction in the absence of STIM1, and the constitutively active L138F mutant channel can cause severe myopathy.^8^ However, the activation mechanism of the L138F Orai1 mutant is not well studied yet. The structure of human Orai1 has not been resolved so far, while the closed state crystal structure of dOrai, which shares 73% sequence identity with human Orai1 within the transmembrane region, has been resolved by Hou et al. at a resolution of 3.35 Å (Fig. 1A).^17^ The L138F mutation in human Orai1 corresponds to the mutation L210F in dOrai of *Drosophila melanogaster*. In order to better understand how L210F mutation activates the channel, molecular dynamics (MD) simulations for the WT and the L210F mutant channels were carried out. Our results revealed a previously undescribed rotation of the residue F210 in the mutant, and the larger rotation angle of F210 in the mutant might be the origin of the activation of the L210F mutant. It was also observed that the rotation of F171 may also play an important role for the activation, as previously reported^27^. Therefore, our simulation results reveal a plausible activation mechanism of the L210F dOrai mutant and may provide a new perspective for understanding the activation mechanism of CRAC channels.

## Materials and Methods

### Molecular dynamics simulations

The crystal structure of *Drosophila melanogaster* Orai obtained from the Protein Data Bank (PDB ID: 4HKR)^17^ was used as the starting structure. MODELLER^28^ was used to build the missing residues of the TM1-TM2 loop (residue number: 181-190) and the TM2-TM3 loop (residue number: 220-235). Both initial simulation systems (Fig. 1B) for the WT dOrai and the L210F dOrai mutant were built using CHARMM-GUI^29^ membrane builder with the channel axis oriented along the z-axis. For each system, the protein was embedded within a 1-palmitoyl-2-oleoyl-sn-glycero-3-Phoss-ph-ocholine (POPC) bilayer with 150 mM NaCl to neutralize the system. The final system size was 110.3 × 110.3 × 105.3 Å^3^ and there were around 116k atoms in the simulation system.

All the molecular dynamics simulations were conducted using GROMACS 5.1.3^30^ with CHARMM36^31–33^ force field and TIP3P^34^ water model. A 500-ps NVT equilibration and a 500-ps NPT equilibration were performed after energy minimization. Then, three 500-ns production simulations were conducted for each system. Position restraints with a force constant of 1000 kJ/mol/nm^2^ were applied on the backbone atoms of the protein for the equilibration simulations. The periodic boundary conditions were used and the time step was 2 fs. The velocity-rescaling algorithm^35^ with a time constant of 0.5 ps was used to maintain the temperature at 310 K. Protein, membrane, and water and ions were coupled separately. The Parrinello-Rahman algorithm^36^ with a time constant of 5 ps was used to maintain the pressure at 1.0 Bar. The Particle-Mesh-Ewald (PME)^37^ method was used to calculate electrostatics and the van der Waals interactions were computed within a cut-off of 1.2 nm. VMD^21^ was used to view trajectories and render figures.

## Results

### Structural statibilty of the WT Orai and the L210F mutant

The root mean square deviation (RMSD) of all six trajectories for both the WT and the L210F mutant were monitored to evaluate their structural changes and stability. The results showed that the systems reached equilibrium states at about 300 ns (Fig. 2A and 2B) with the C*α*-RMSD of the WT and the mutant converging to 0.41 nm and 0.47 nm respectively. This time scale and the RMSD values were slightly larger than the work of Amcheslavsky et al.^38^ and Frischauf et al.^39^, in which the values were roughly 120 ns and 0.2 nm. Our results were more comparable to the work of Dong et al.^25^ owing to similar simulation setups. In addition, the RMSD of the mutant experienced a larger increase during the initial stage of the simulations, reflecting the influence of the residue mutation on the structure (Fig. 2B).

**Figure 2.**
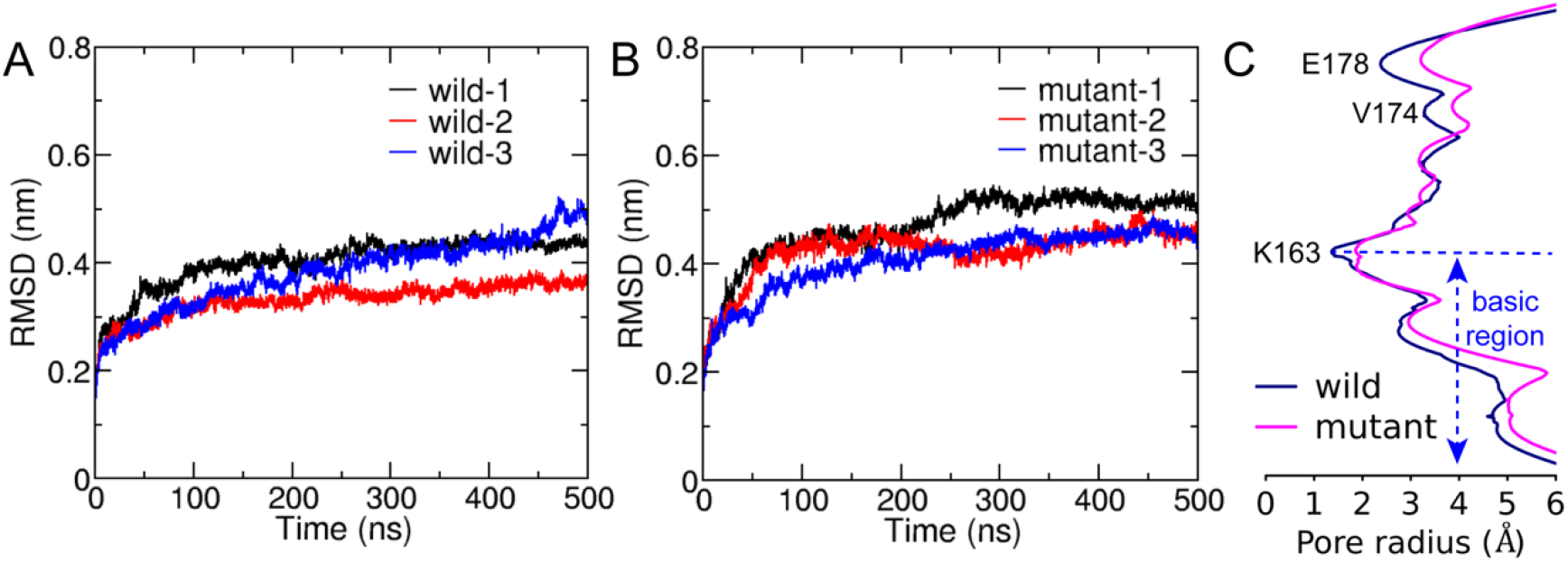
The structural stability during simulations and the average pore radius after reaching equalibrium. C*α*-RMSD of the WT (A) and the L210F mutant (B). Three trajectories are represented in red, blue and black, respectively. (C) The average pore radii. The average pore radii for the WT and the L210F mutant were obtained by calculating the pore radius of the average structures obtained from the last 200 ns of all of the three trajectories for each protein. Hole 2.0^40^ was used to calculate the radius with all the hydrogen atoms removed in the calculation.

The pore radius was calculated to evaluate the effect of the L210F mutation on the channel state. The simulation results revealed three significant radius changes of the mutant (Fig. 2C). The first one occurred in the SF, the glutamate ring - E178, which binds and transports Ca^2+^ selectively. The pore radius at E178 of the L210F mutant expanded by approximately 1 Å compared with the WT channel. An increase in the radius of the SF may increase the chance of ion binding and thus improve the probability of ions entering into the pore for the mutant. The second change was at the starting residue of the hydrophobic region, V174. The dilation of V174 in the mutant may allow more water molecules to stay at this entrance of the hydrophobic region, which lays a good foundation for water to further occupy the following hydrophobic region. The third change occurred in the basic region, where nearly the whole segment of the mutant was wider than the WT. The increase of the basic region radius can not only reduce the steric hindrance, but also reduce the electrostatic exclusion between the basic residues and cation ions passing through this cationic channel. Moreover, the constriction site of the channel, the K163 gate located at the beginning of the basic region, expanded significantly in the mutant channel (Fig. 2C), which might be a key step for the activation of the L210F mutant.

### The rotation of the residue L/F210

Two carbon atoms of L/F210 in the starting structure of the simulation were selected to define the initial vector (Fig. 3A). The rotation angle of L/F210 in each frame was then defined as the angle between the initial vector and the vector linking the two same carbon atoms in each frame of the simulation (Fig. 3A), which was used to evaluate the conformational changes of L/F210 in the WT and L210F mutant channels. The results revealed a previously undescribed rotation of the residue L/F210. F210 in the mutant channel exhibited less distribution of rotations that are less than 20 degrees, while plenty of distribution of rotations that are larger than 77 degrees, which is distinct from L210 in the WT (Fig. 3B). The more distributed F210 with larger rotation angles, especially those larger than 77 degrees, kept this residue farther away from the pore-lining helix (TM1) (Fig. 3A), which will probably generate a pulling effect on the TM1 through hydrophobic interactions with A166 on the TM1 of the same subunit (Fig. S2) and leave more room for the TM1 to expand outward. This may be the reason that caused the expansion of the basic region located in the TM1 adjacent to F210 in the mutant channel (Fig. S2).

**Figure 3.**
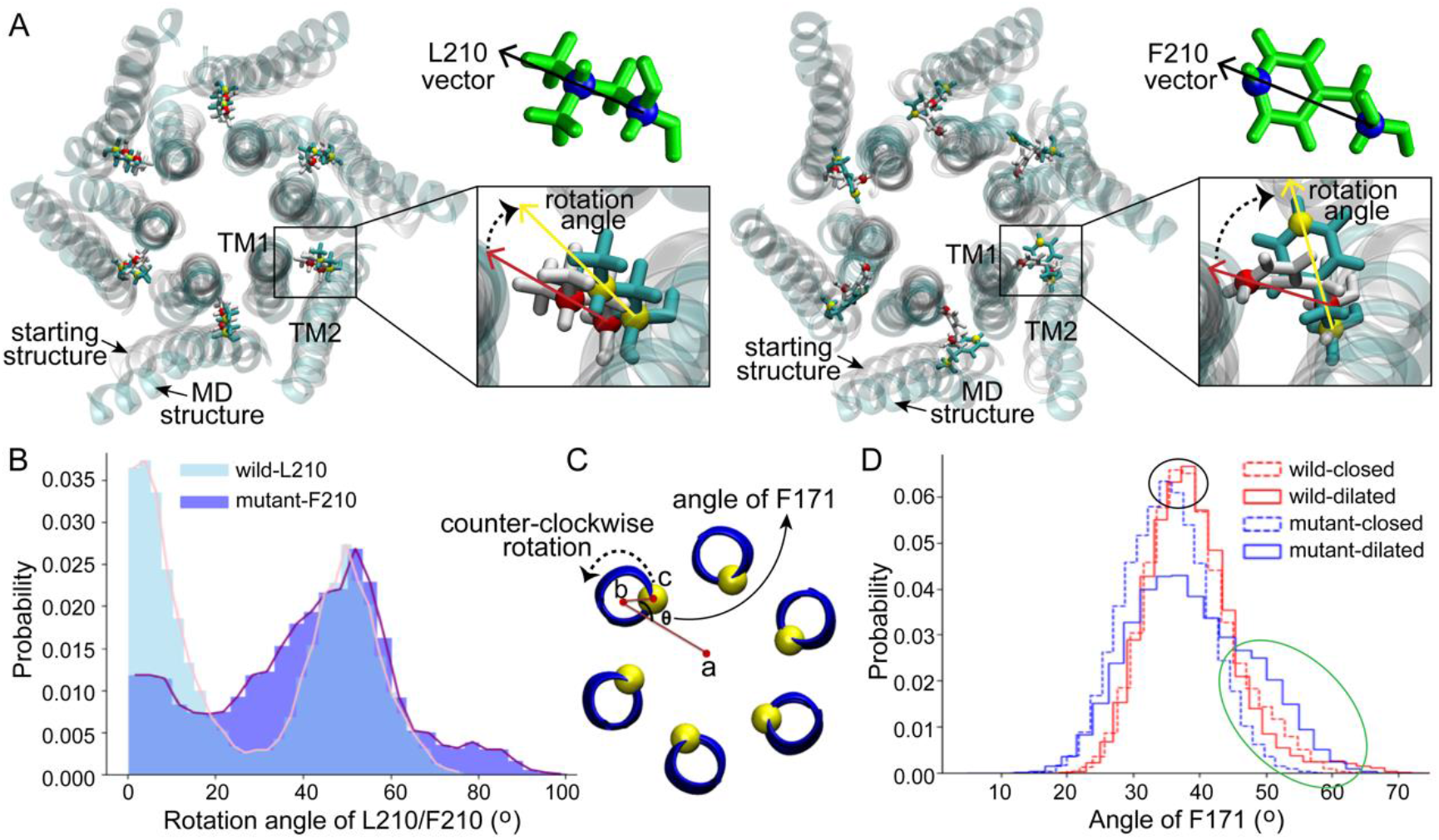
The rotation of residues L/F210 and F171. (A) The definition of the rotation angle of L/F210. Only the TM1 and TM2 in each subunit are shown for clarity. The starting structure of the simulation is shown with white transparent NewCartoon, and one frame of the simulation is shown with cyan transparent NewCartoon as an example. L210 and F210 are shown with Licorice. (B) The distribution of the rotation angle of L/F210. The last 200 ns of each trajectory was used for this analysis. (C) The definition of the angle of F171. Point a was obtained by projecting the center of the channel on the XY plane. Similarly, point b was determined by the mass center of the two helices centered on F171, and point c was the center of the C-α of F171. (D) The distribution of the angle of F171. The four data sets, wild-closed, wild-dilated, mutant-closed and mutant-dilated, were classified with a pore radius of 2 Å at the K163 gate. 500 ns of each trajectory was used for the analysis.

### The counter-clockwise rotation of the residue F171

It was reported that the opening of the Orai channel is accompanied by the counter-clockwise rotation of the residue F171,^27^ which is located on the TM1 in the middle of the hydrophobic gate and is some distance away from the mutation point L/F210 (Fig. 1A and Fig. S2). The presence of the hydrophobic gate increases the energy barrier of ion permeation, while the rotation of the residue F171 may reduce this barrier, contributing to the activation of the channel^27^. Here, the orientation angle of F171 (definition in Fig. 3C) was calculated to measure the dynamics of this residue in our simulations. Firstly, in order to investigate the angle of F171 in a more detailed pore radius range, the structures of each channel obtained from MD simulations were classified into two classes according to the pore radius at the K163 gate, which is the constriction site of the channels. Structures with a radius at K163 of less than 2 Å were classified as the closed state while structures with a K163 radius of more than 2 Å were classified as the dilated state. As a result, four data sets: wild-closed (10123 frames), wild-dilated (4877 frames), mutant-closed (5175 frames) and mutant-dilated (9825 frames) were obtained. The results of the angle showed a very similar distribution among the wild-closed, wild-dilated and mutant-closed data sets with the most frequent angles of 35-40 degrees (Fig. 3D, the black oval). These distributions were supposed to be caused by the normal fluctuations of F171. However, the angle distribution of the mutant-dilated data set was different, with a probability increase of angles above 45 degrees (Fig. 3D, the green oval, about 45~65 degrees) and a probability decrease of angles around 35~40 degrees. A larger rotation angle of F171 will keep this residue pointing away from the pore axis, which will further allow more hydration at this site. This appears to be caused by the dilation of V174, on the basis of the structural change at F210 in the mutant (Fig. 2C), and will lead to a more open hydrophobic gate. Therefore, the angles above 45 degrees were believed to make contributions to the opening of the hydrophobic gate of the mutant. Then, the effective counter-clockwise rotation of F171 that might open the hydrophobic gate was about 10-30 degrees (45\65 minus 35) after eliminating its normal fluctuations. The average value of 20 degrees is similar to the counter-clockwise rotation of 19 and 20 degrees in another two constitutively open channels V174A and F171Y.^27^

### Water in the pore

The Orai channel has two gates, the residue K163 gate in the basic region and the hydrophobic gate (residues F171-V174). Previous studies have shown that even if the radius of the Orai channel does not significantly change, a limited increase of hydration in the pore is enough to regulate the conduction state^25^, suggesting the importance of the hydrophobic gate. Hence, the number of water molecules in the pore, as an important indicator of the conductivity of the channel, was calculated to measure the hydration difference between the WT and the L210F mutant channels. The water distributions in the region from residues F171 to E178 were measured. This region includes the hydrophobic region and the SF, which was reported to be the area of the most significant water distribution difference.^25^ The results showed that the distribution of the number of water molecules was relatively concentrated with the average number of 16 in the pore of the WT (Fig. 4A). However, the distribution showed two peaks in the L210F mutant (Fig. 4B). A low-hydration distribution and a high-hydration distribution were observed, and the average number for the whole distribution was about 18 (Fig. 4B), larger than the average number in the WT. Besides, the mutant can have a hydration number as high as ~30, a value the WT channel never reached. Accordingly, the highly hydrated structures of the L210F mutant were supposed to reduce the energy barrier of ions passing through the hydrophobic gate and thus facilitate ion permeation.

**Figure 4.**
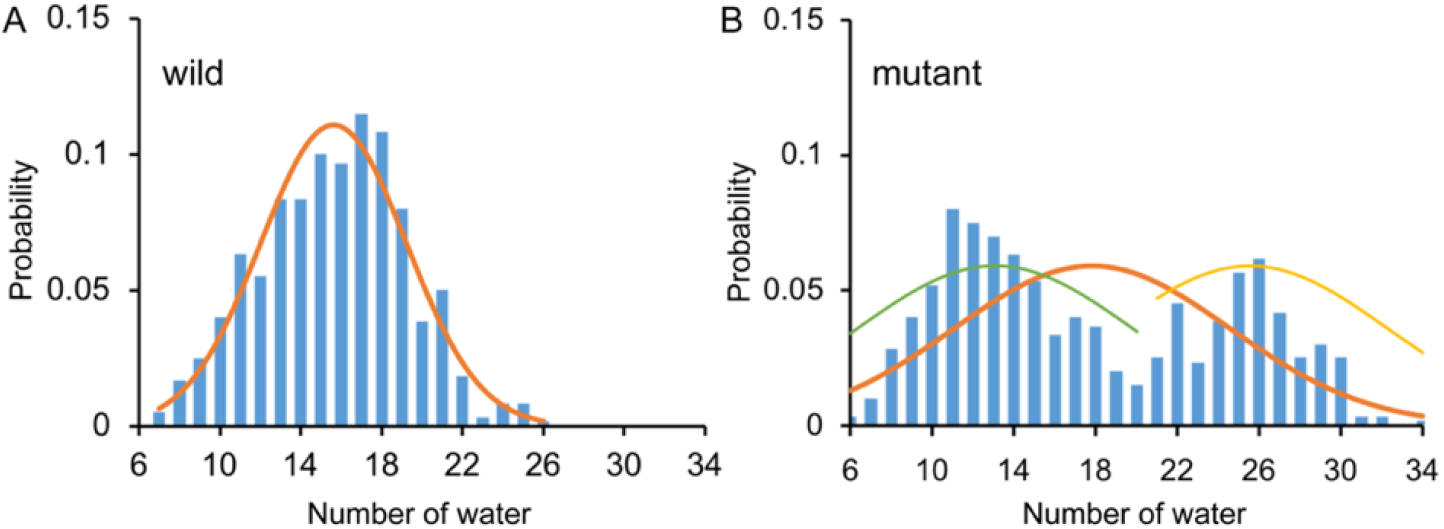
The number of water molecules and the corresponding probability in the pore of the WT (A) and the L210F mutant (B) channels. From residue F171 to E178 along the central axis of the pore, the number of water molecules (oxygen atoms were measured) within a cylinder of 5-Å radius was calculated using VMD for the last 200 ns of all three trajectories for each protein system.

### Na^+^ in the pore

Although the Orai channels are highly selective for Ca^2+^, Na^+^ was often used in the study of the conductivity of CRAC channels as Na^+^ can permeate at a much higher rate in the absence of Ca^2+^,^15, 16^ which makes it easier for better sampling in MD simulations. The position and the number of Na^+^ ions within the pore were calculated to observe the behavior of Na^+^ ions in the WT and the L210F mutant channels. The results showed that Na^+^ ions were mainly distributed in the upper part of the pore (Fig. 5A and 5B). Three binding sites, residues E178, D182 and D184, were observed (Fig. S1), among which E178 was the most dominant one. Na^+^ ions permeated deeper in the mutant than in the WT (below E178 in the mutant and a little above E178 in the WT, Fig. 5A and 5B), and the most frequent numbers of Na^+^ ions in this region were 7 and 9 for the WT and the mutant, respectively (Fig. 5C and 5D). Therefore, it seems that the L210F mutation moderately modified the distribution of Na^+^ ions along the pore, causing more Na^+^ accumulation at the entrance of the pore and increasing the probability of ions passing through, which is consistent with the fact that the L210F mutant is constitutively open to cations. However, no spontaneous Na^+^ permeation was observed in our MD process, probably due to the extremely low conductivity of the L138F Orai1 (equals to the L210F dOrai here) mutant^39^ and the lacking of a fully open structure.

**Figure 5.**
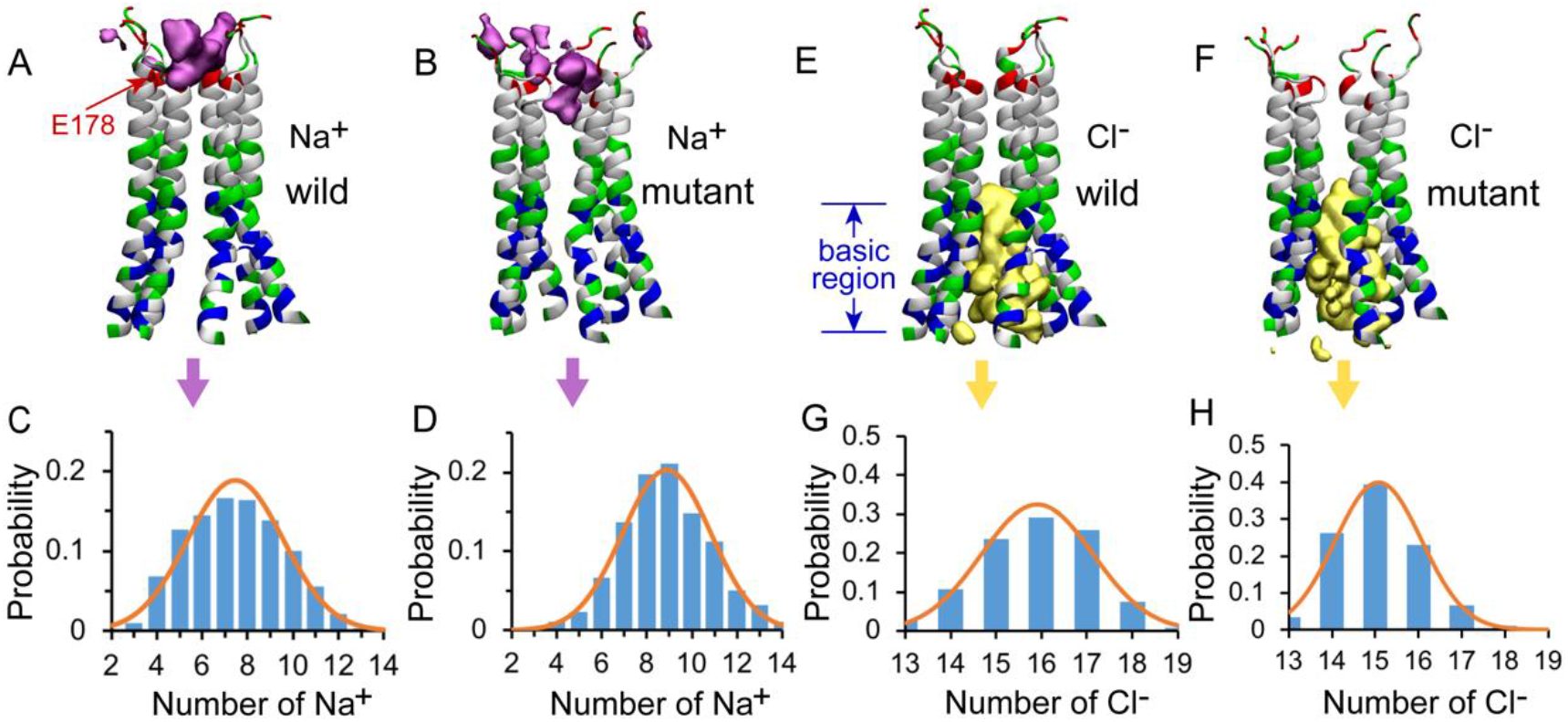
The distributions of Na^+^ and Cl^−^ in the pore of the channel. The isosurface of Na^+^ ion density in the WT (A) and the L210F mutant (B) channels. The number of Na^+^ ions and corresponding probability in the pore of the WT (C) and the L210F mutant (D) channels. The isosurface of Cl^−^ ion density in the WT (E) and the L210F mutant (F) channels. The number of Cl^−^ ions and corresponding probability in the pore of the WT (G) and the L210F mutant (H) channels. The isosurface of Na^+^ and Cl^−^ ion densities are shown in pink and yellow with isosurface values of 0.02 and 0.06, respectively. Only residues from W148 to D184 are shown with NewCartoon and colored by residue types for clarity. Blue, basic residues; red, acidic residues; green, polar residues; white, nonpolar residues. From residue W148 to D184 along the central axis of the pore, the number of Na^+^ and Cl^−^ ions within a cylinder of 10-Å radius were calculated using VMD for the last 200 ns of all three trajectories for each protein system.

### Cl^−^ in the pore

As previous studies showed anion-assisted cation permeation in the V174A Orai channel (also a constitutively open channel),^41^ the position and number of Cl^−^ ions within the pore were calculated to analyze the behavior of Cl^−^ ions in the WT and the L210F mutant channels in our simulations as well. The results showed that Cl^−^ ions were mainly distributed in the intracellular region, specifically referring to the basic region including residue K163 and the residues below it (Fig. 5E and 5F). The most frequent numbers of Cl^−^ ions were 16 and 15 in the WT and the mutant channels, respectively (Fig. 5G and 5H), showing no appreciable difference. Previous molecular dynamics simulations reported that Cl^−^ ions in the basic region of the open V174A mutant can flow out to the extracellular side of the pore under electric field conditions, coordinating with Na^+^ ions in the pore to form an energetically more favorable cluster to help the influx of Na^+^ ions.^41^ It seems that the mutation L210F does not alter the Cl^−^ occupation around the basic region or its role in assisting cation permeation.

## Discussion

In this paper, based on the crystal structure of *Drosophila melanogaster* Orai, we investigated the detailed channel structures and the water and ion distributions for both the WT and the L210F mutant channels by using molecular dynamics simulations. The results revealed two small but essential conformational changes resulted from the L210F mutation. Firstly, an undescribed rotation of residue F210 initiated the channel opening in the L210F mutant. F210 in the mutant had larger outward rotation than L210 in the WT, resulting in the dilation of the basic region and the K163 gate. At the same time, a 20-degree (on average) counter-clockwise rotation of F171 in the hydrophobic gate occurred more frequently and allowed more hydration at the hydrophobic region, which can potentially lead to the opening of the hydrophobic gate. Collectively, the rotation of F210 and F171, leading to the opening of the two gates of the channel, may create a constitutively open L210F mutant. Therefore, our results may shed further light on the disease of myopathy caused by the L138F mutation in human Orai1, by providing insight into the detailed structure and activation mechanism of the L210F mutant.

This is the first time that the rotation of the residue 210 is characterized to be the key origin of the activation for the L210F mutant channel. Compared with L210 in WT, the larger rotation angle of F210 induced the expansion of the basic region and the K163 gate. The expansion of the basic region involved hydrophobic interactions between TM2 and TM1 in the same subunit, suggesting the important role of the transmembrane helix (TH) interaction network on the channel gating. Apparently, the regulation of the TH network may work in more than one way. In Frischauf’s work for another constitutively open H134A Orai1 mutant (equals to the H206A dOrai mutant), the regulation of the TH connectivity on the channel gating is shown in the disruption of hydrogen bonds between H134 on the TM2 and two residues on the TM1 (S93 and S97).^39^ Notably, the L138F Orai1 mutant (equals to the L210F dOrai here) is also studied through molecular dynamics simulations in the same work, which showed that the enhanced hydrophobic contacts between TM2 and TM1 through L138F mutation trigger more flexibility of the pore, and then one water chain enters into the hydrophobic region and opens the pore.^39^ These results are generally consistent with ours, both of which emphasize the importance of the hydrophobic interaction between TM2 and TM1 and water chain or hydration in the hydrophobic region. However, our results revealed a previously unnoticed rotation of residue 210, which may be the origin of the activation. Apart from the rotation of the residue 210, the rotation of F171 observed in our simulation, which was also reported to be required in other constitutively open channels V174A and F171Y in Yamashita’s work^27^, may also be an important factor for gating.

Notably, no significant rotational movement of F171 or TM1 was observed from the open conformation of the H206A dOrai (equals to the H134A Orai1) resolved at 3.3 Å resolution by cryo-EM recently^42^, indicating that multiple activation mechanisms may be utilized by different mutants. The regulation of the TH network is also shown in the dilation of the filter. TM1, TM2 and TM3 in two adjacent subunits may participate in this process (Fig. S3). The residue K270 on the TM3 was reported to regulate the filter selectivity through conformation dynamics and the filter dilation was also observed in the previous study.^43^ In our study, the radius of the K270 ring was dilated in the L210F mutant (the average radius of K270 ring in the WT and the mutant: 12.30 Å and 13.33 Å), which was probably caused by the expansion of the filter through electrostatic interactions (Fig. S3). However, how the mutation of residue 210 allosterically affects K270 is still not clear. Further simulations and analysis are undergoing to focus on the selectivity difference between the WT and the L210F mutant using a new calcium model,^44^ which will hopefully provide a better understanding on this aspect.

The dilation of the entire pore obtained in our simulations for the L210F mutant is consistent with the recently obtained open-state cryo-EM structure of the H206A dOrai mutant^42^. Both H206A and L210F mutations occur on the TM2 and construct constitutively open channels with some selectivity for Ca^2+^ remained, so some structural similarity of the pore may exist among the H206A mutant, the L210F mutant and the WT Orai.^39^ The dilation of the filter in the open channels didn’t get much attention previously. In some previous simulations^39^, the filter of the constitutively open channel is nearly the same with the closed Orai, since Ca^2+^ were trapped in the filter in both simulations owing to strong interactions between Ca^2+^ and protein. Here, we didn’t include Ca^2+^ in our simulations, following the protocol of other previous simulations^25^, and therefore the filter was more flexible to reach a more dilated state in the mutant, in agreement with the filter dilation as observed in the open-state H206A structure^42^. In addition, it was reported that the WT Orai1 experiences some filter conformational changes when it is activated by STIM1, suggesting that the filter conformation may be different for the open channel and the closed channel.^24^ Still, it should be noted that both the L210F Orai mutant and the H206A cyro-EM structure may not fully mimic the STIM-dependent Orai gating since their conductivity and selectivity are not the same after all.

## Acknowledgements

We thank Xiaolan XU who inspired us to work on this project. The research was supported by the National Natural Science Foundation of China (21873006 and 32071251 to CS), and the National Key Research & Development Program of the Ministry of Science and Technology of China (2016YFA0500401 and 2021YFE0108100 to CS). Part of the molecular dynamics simulation was performed on the Computing Platform of the Center for Life Sciences at Peking University.

## Supporting Information

**Figure S1.**
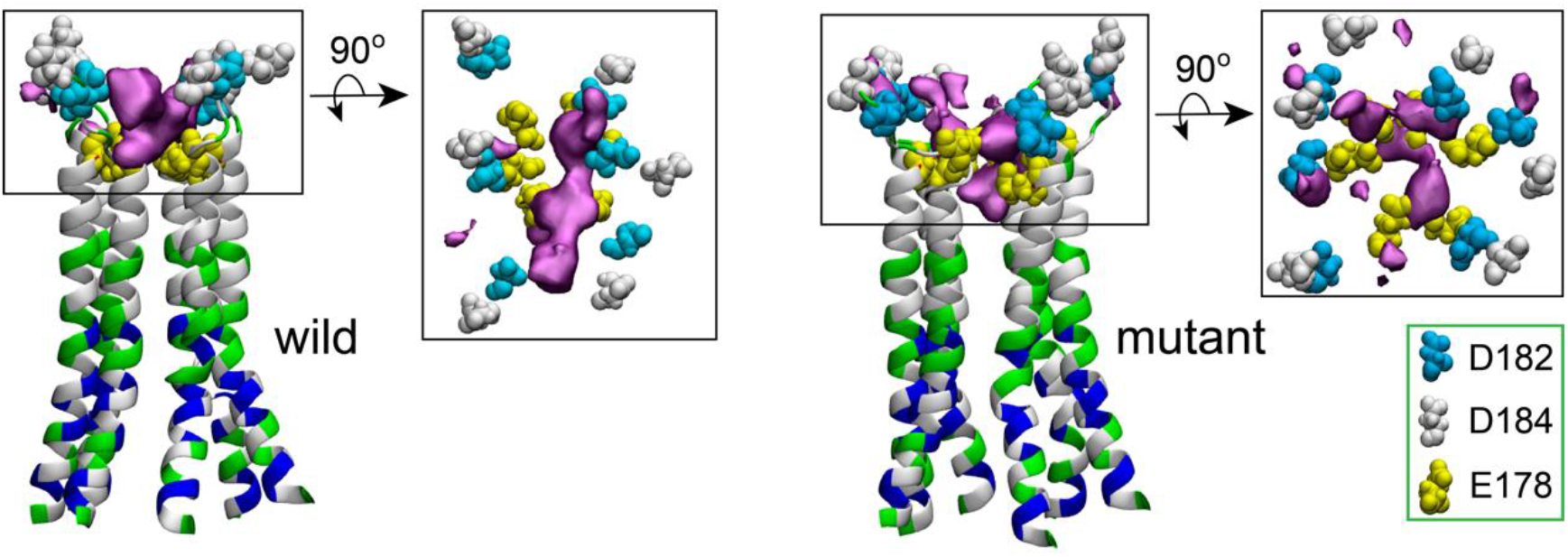
The Na^+^ binding sites. The three binding sites E178, D182 and D184 are shown with VDW spheres. The isosurface of Na^+^ ion densities are shown in pink with isosurface value of 0.02. Only residues from W148 to D184 are shown in NewCartoon and colored by residue types for clarity. Blue, basic residues; red, acidic residues; green, polar residues; white, nonpolar residues.

**Figure S2.**
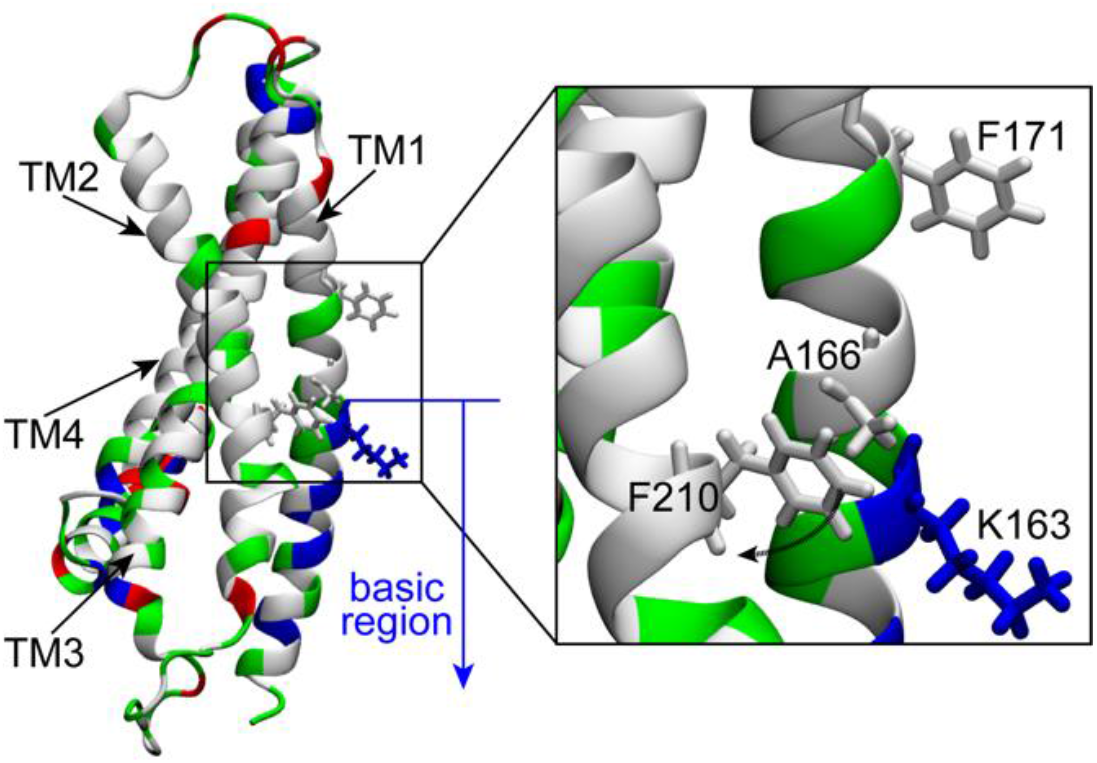
The location of F210, A166, K163 and F171. The outward rotation of F210 might be the reason of the dilation of the basic region. Only one subunit is shown for clarity.

**Figure S3.**
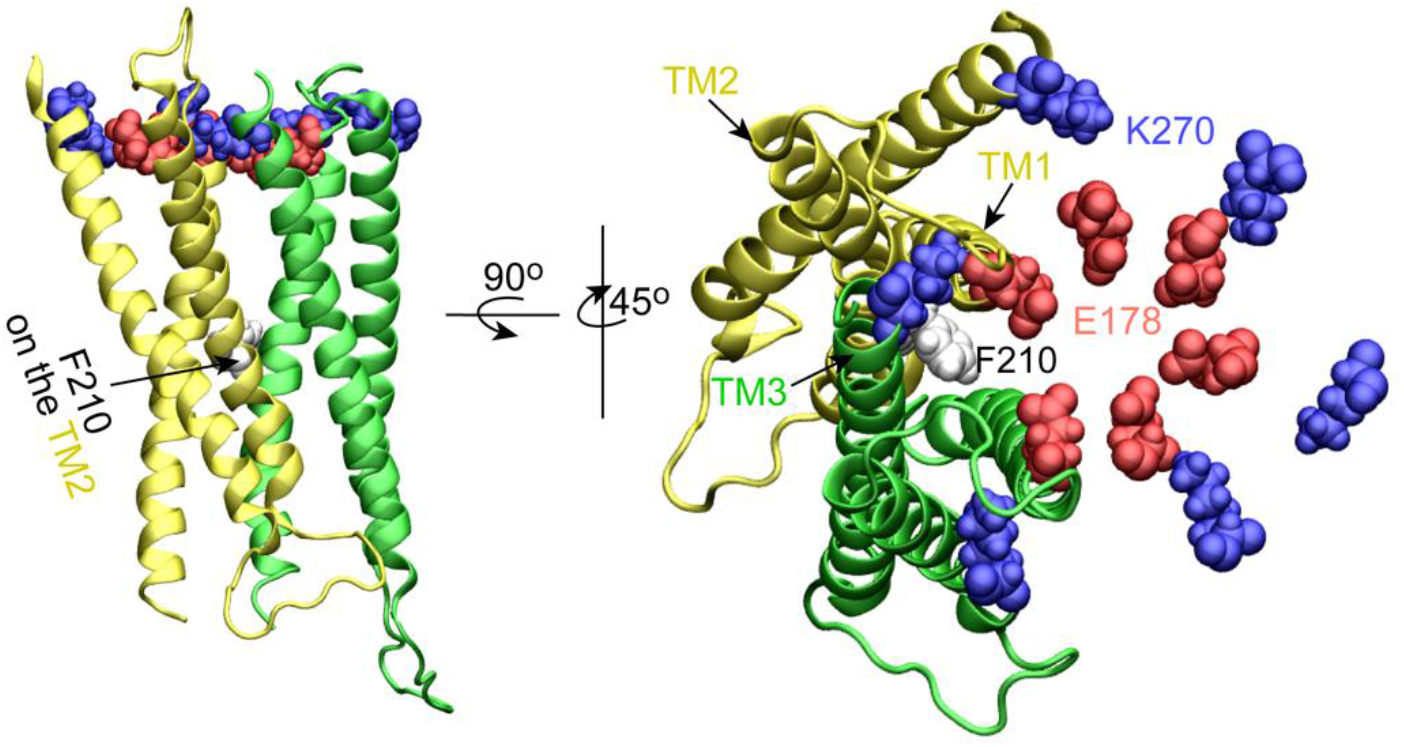
The location of F210 and the dilation of the SF. Two subunits are shown with yellow and green NewCartoon, respectively. Only the TM1, TM2 and TM3 are shown for clarity. F210 on the TM2 is shown with white VDW spheres. K270 on TM3 and E178 (the SF) on TM1 in all of the six subunits are shown with blue and red VDW spheres, respectively.

